# A Ketogenic Diet Sensitizes Pancreatic Cancer to Inhibition of Glutamine Metabolism

**DOI:** 10.1101/2024.07.19.604377

**Authors:** Omid Hajihassani, Mehrdad Zarei, Asael Roichman, Alexander Loftus, Christina S. Boutros, Jonathan Hue, Parnian Naji, Jacob Boyer, Soubhi Tahan, Peter Gallagher, William Beegan, James Choi, Shihong Lei, Christine Kim, Moeez Rathore, Faith Nakazzi, Ishan Shah, Kevin Lebo, Helen Cheng, Anusha Mudigonda, Sydney Alibeckoff, Karen Ji, Hallie Graor, Masaru Miyagi, Ali Vaziri-Gohar, Henri Brunengraber, Rui Wang, Peder J. Lund, Luke D. Rothermel, Joshua D. Rabinowitz, Jordan M. Winter

## Abstract

Pancreatic cancer is the third leading cause of cancer death in the United States, and while conventional chemotherapy remains the standard treatment, responses are poor. Safe and alternative therapeutic strategies are urgently needed^1^. A ketogenic diet has been shown to have anti-tumor effects across diverse cancer types but will unlikely have a significant effect alone. However, the diet shifts metabolism in tumors to create new vulnerabilities that can be targeted (1). Modulators of glutamine metabolism have shown promise in pre-clinical models but have failed to have a marked impact against cancer in the clinic. We show that a ketogenic diet increases TCA and glutamine-associated metabolites in murine pancreatic cancer models and under metabolic conditions that simulate a ketogenic diet *in vitro.* The metabolic shift leads to increased reliance on glutamine-mediated anaplerosis to compensate for low glucose abundance associated with a ketogenic diet. As a result, glutamine metabolism inhibitors, such as DON and CB839 in combination with a ketogenic diet had robust anti-cancer effects. These findings provide rationale to study the use of a ketogenic diet with glutamine targeted therapies in a clinical context.

**Graphical Abstract:** 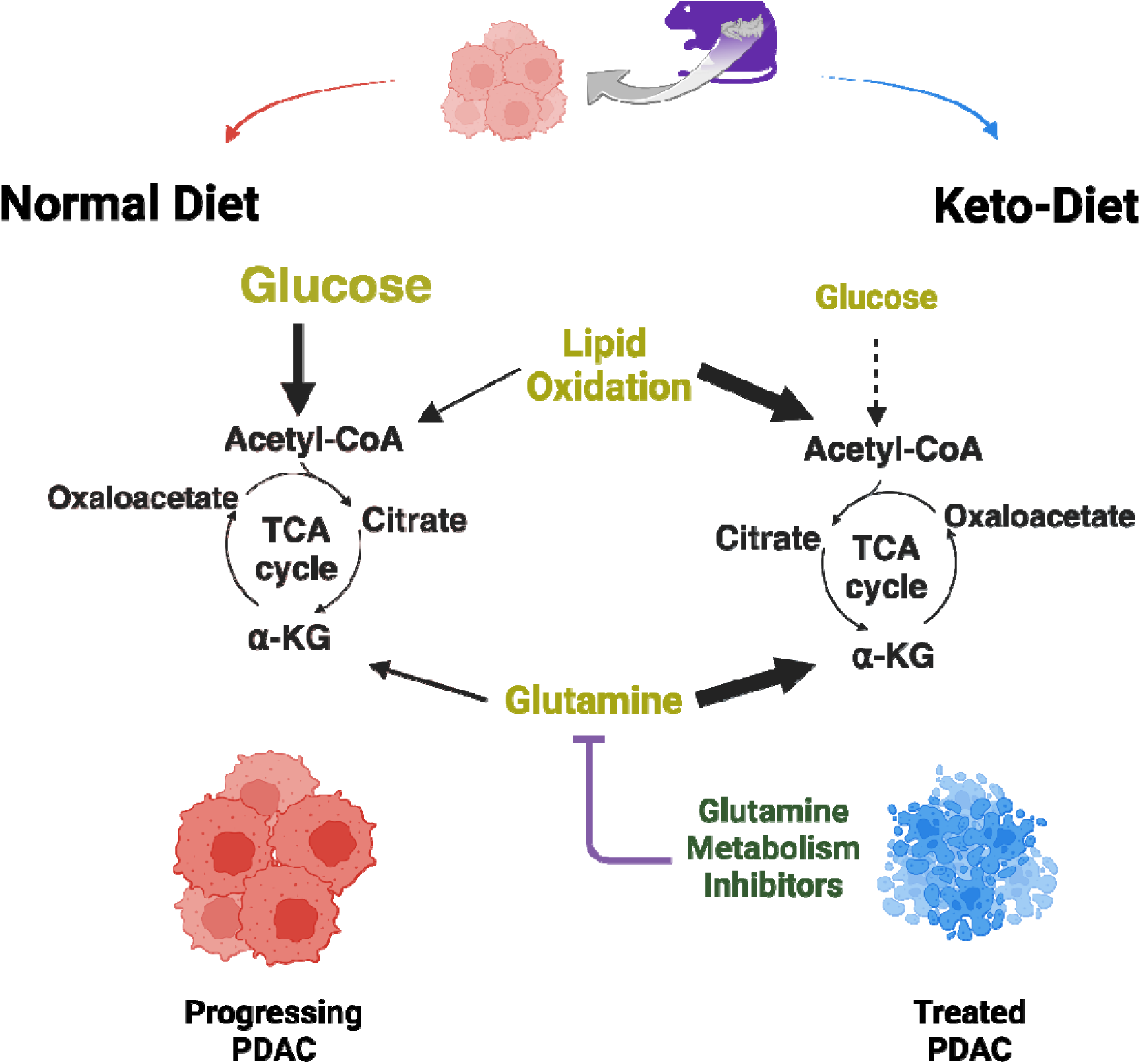

Graphical Abstract Description: **Mechanistic rationale for combining a ketogenic diet and glutamine metabolism inhibitors.** The combination of low glucose from a ketogenic diet and pharmacologic glutamine inhibition impairs nutrient input to mitochondria, reducing cancer growth.

## Introduction

Pancreatic ductal adenocarcinoma (PDAC) continues to have a poor prognosis, with a 5-year overall survival rate of approximately 12% (2). A complex and dense tumor microenvironment (TME) is believed to be a prominent contributor to PDAC aggressive biology (3–5). A haphazard microvasculature and associated steep nutrient gradients activate survival pathways in PDAC cells, which drive treatment resistance and promote cancer progression (6). Numerous studies have shown that PDAC cells adapt to a challenging and nutrient limited TME by reprogramming metabolism towards more efficient energy extraction. This metabolic shift involves a greater utilization oxidative phosphorylation by mitochondria (7). Alternative energy substrates beyond glucose are required to feed the tricarboxylic acid (TCA) cycle since glucose is typically limiting in tumors (8), and these alternative nutrients can be sourced through diverse adaptive processes like autophagy, macropinocytosis, and anaplerosis of non-glucose organic intermediates (9). Despite significant advancements in the understanding of PDAC metabolism over the past decade, translation of this knowledge into clinically applicable and effective therapeutic strategies has eluded researchers.

A ketogenic diet, alone or in combination with other therapies has displayed an anti-cancer signal across a wide array of cancer subtypes in clinical and pre-clinical studies (10). While dietary modification alone probably will not impact PDAC survival in a clinically meaningful way, the physiological changes induced by a ketogenic diet, particularly within a nutrient-limited PDAC TME, may push conditions to a point that creates exaggerated metabolic dependencies, exposing actionable therapeutic vulnerabilities in cancer cells but not in normal tissues. Thus, insights into the metabolic changes induced by a ketogenic diet could nominate specific therapeutic partners to potentiate anti-tumor effects of a ketogenic diet (1,11,12). For instance, a ketogenic diet has been shown to increase circulating glutamine and related metabolites in animal models (13). Glutamine (Gln) plays a multifaceted role in cellular metabolism. The organic backbone contributes to the generation of TCA cycle intermediates through anaplerosis via α-ketoglutarate (14–16), and the amide nitrogen fuels the biosynthesis of essential metabolites like asparagine and nucleotides (17,18).

Previous work by our group revealed that PDAC cells preferentially metabolize Gln when glucose levels are reduced (19). Since one of the key impacts of a ketogenic diet is to lower systemic and intra-tumoral glucose levels, due to substantially reduced carbohydrate intake (12), we reasoned that this feature of a ketogenic diet might enhance reliance on glutamine metabolism as an alternative fuel source in pancreatic cancer cells (20,21). Glutaminase (GLS) (which converts glutamine to glutamate prior to entry into the TCA cycle) inhibition, has been explored in pancreatic cancer models as a target of single-agent therapy, and the treatment transiently reduced cellular proliferation *in vitro*. However, this approach was ineffective across multiple *in vivo* PDAC models as a monotherapy due to compensatory metabolic reprogramming (17). It was postulated that broadening the inhibition of Gln metabolism to related targets beyond GLS could enhance efficacy and thwart adaptive rewiring changes responsible for resistance. For instance, a Gln analog, 6-diazo-5-oxo-1-norleucine (DON), which covalently and irreversibly binds to multiple Gln-metabolizing enzymes, broadly impedes Gln metabolism to a superior degree, including its utilization in the generation of hexosamines and nucleotides (14). These studies hint at the superiority of a less selective pharmacologic approach.

Herein, we explore whether a ketogenic diet exposes a vulnerability to glutamine metabolism inhibition through induction of a low glucose state in PDAC. Additionally, we attempt to illuminate how lipids and ketone bodies supplied by a ketogenic diet are utilized by cancer cells to compensate for low glucose and low protein levels, which could expose additional metabolic targets. This line of investigation offers an example of a “push-pulse” strategy, where a ketogenic diet nudges a system towards greater dependency on a specific metabolic program, which in turn exposes new dependencies (22).

## Results

### A ketogenic diet inhibits tumor progression in mice

We tested the effects of ketogenic diet alone, compared to a control diet, in a series of experiments involving MIA-PaCa2 (derived from human PDAC) xenografts or orthotopic KPC (derived from murine PDAC) xenografts in nude mice. In these experiments, tumor measurements and dietary intervention commenced when a palpable tumor size of approximately 150 mm² was reached after two consecutive tumor growth measurement. The control diet contained 7% calories from fat (AIN-93G), while the ketogenic diet comprised 90% of calories from fat. We observed a robust suppression of PDAC growth associated with the ketogenic diet alone across both models (Fig. 1 A, B; Supplementary Fig.1 A). To our surprise, this ketogenic diet induced anti-tumor effect was reproducibly stronger than previous observations by others (23,24). Mice on the ketogenic diet exhibited notable reductions in circulating glucose levels, maintaining a range of 100-160 mg/dl, compared to mice on the standard diet where glucose levels ranged from 140 to 220 mg/dL (Fig 1C and Supplementary Data Fig. 1B). We confirmed ketosis with a ketogenic diet through measurements of circulating β -hydroxybutyrate levels (mean of 1.5 vs. mean of 0.5 in controls, Fig. 1C). While published studies by other suggest have shown a sharp drop in body weight when transitioning to a ketogenic diet (25), we observed an initial drop of 10-15% with a ketogenic diet, followed by weight stability beyond day 15 of the study (Fig. 1D, Supplementary Data fig 1C).

**Figure 1:**
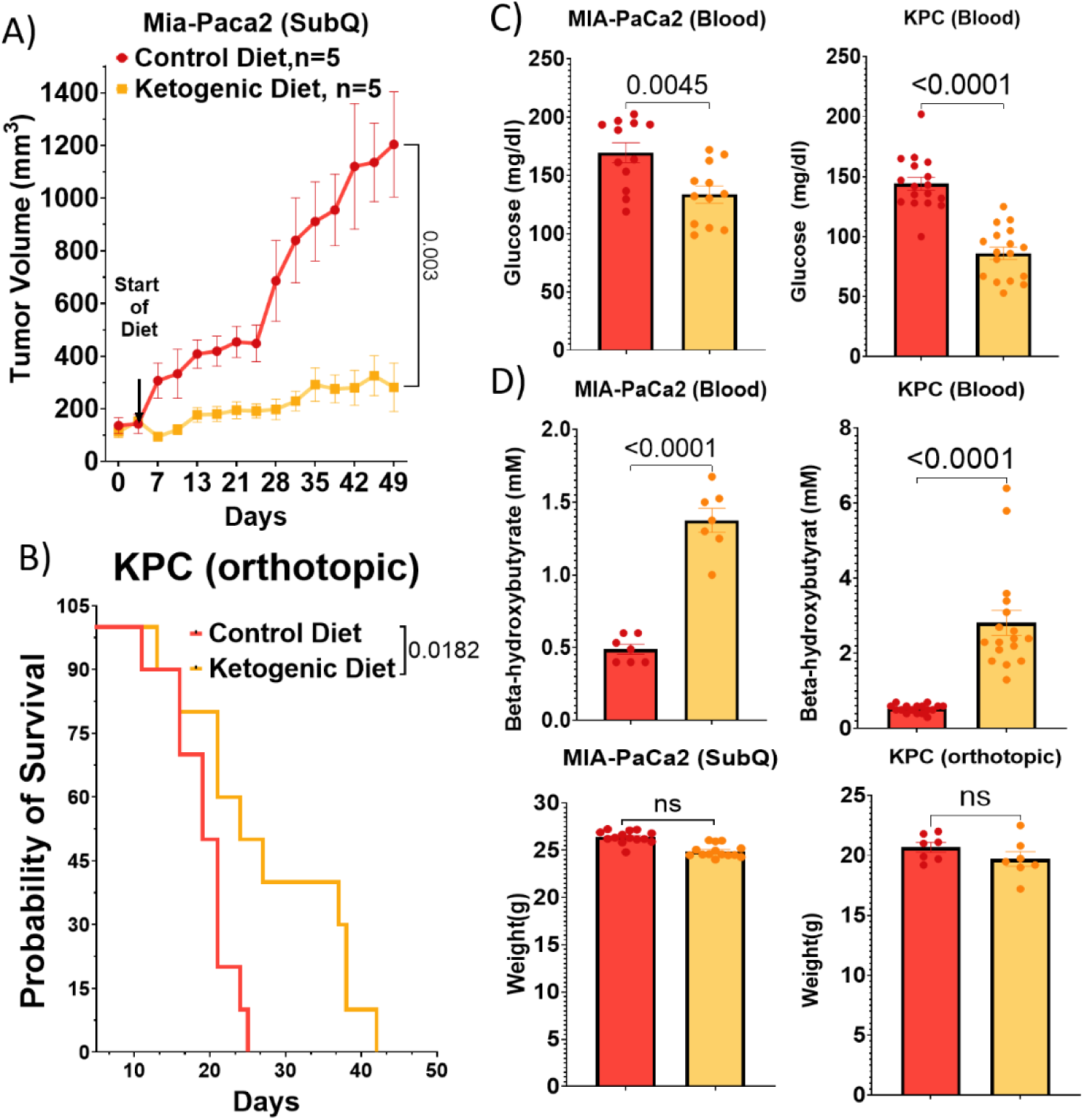
Suppression of Tumor Growth in PDAC Models By a Ketogenic Diet. **(A)** Subcutaneous injection of MIA-PaCa2 cells in athymic nude mice (n=5). **(B)** Survival probability after orthotopic injection of KPC cells into the pancreas of C57BL/6 mice (n=10). **(C)** Levels of glucose and of β-hydroxybutyric acid in blood of C57BL/6 and athymic nude mice undergoing a normal diet and a ketogenic diet (n=5). **(D)** Body weights of treated mice compared to the normal diet

### Enhanced glutamine and TCA cycle metabolite levels in serum and tumors exposed to a ketogenic diet

A ketogenic diet reprograms the metabolic landscape of PDAC tumors and changes are detectable in the host serum in mice. The dietary intervention resulted in elevated intratumoral glutamine levels in MIA-PaCa2 tumors (Fig. 2A). Correspondingly, intratumoral levels of glutamate (derived from glutamine by GLS), aspartate (generated from glutamate by GOT), ketone bodies, and TCA cycle metabolites (potentially fueled by glutamine-glutamate anaplerosis) were all increased (Fig 2A & B). Notably, changes in intratumoral glutamine mirrored those observed in circulating glutamine levels (Fig. 2D). Similarly, circulating ketone bodies in the serum correlated with intratumoral β-hydroxybutyric acid concentrations (Fig 2E). Furthermore, altered fatty acid profiles within tumors and serum reflect the diet’s impact on circulating fatty acids (Fig 2C & F). Collectively, these findings suggest a direct impact of a ketogenic diet on PDAC tumor glutamine metabolism, potentially fueling key metabolic functions like oxidative phosphorylation in the context of diminished circulating glucose levels.

**Figure 2:**
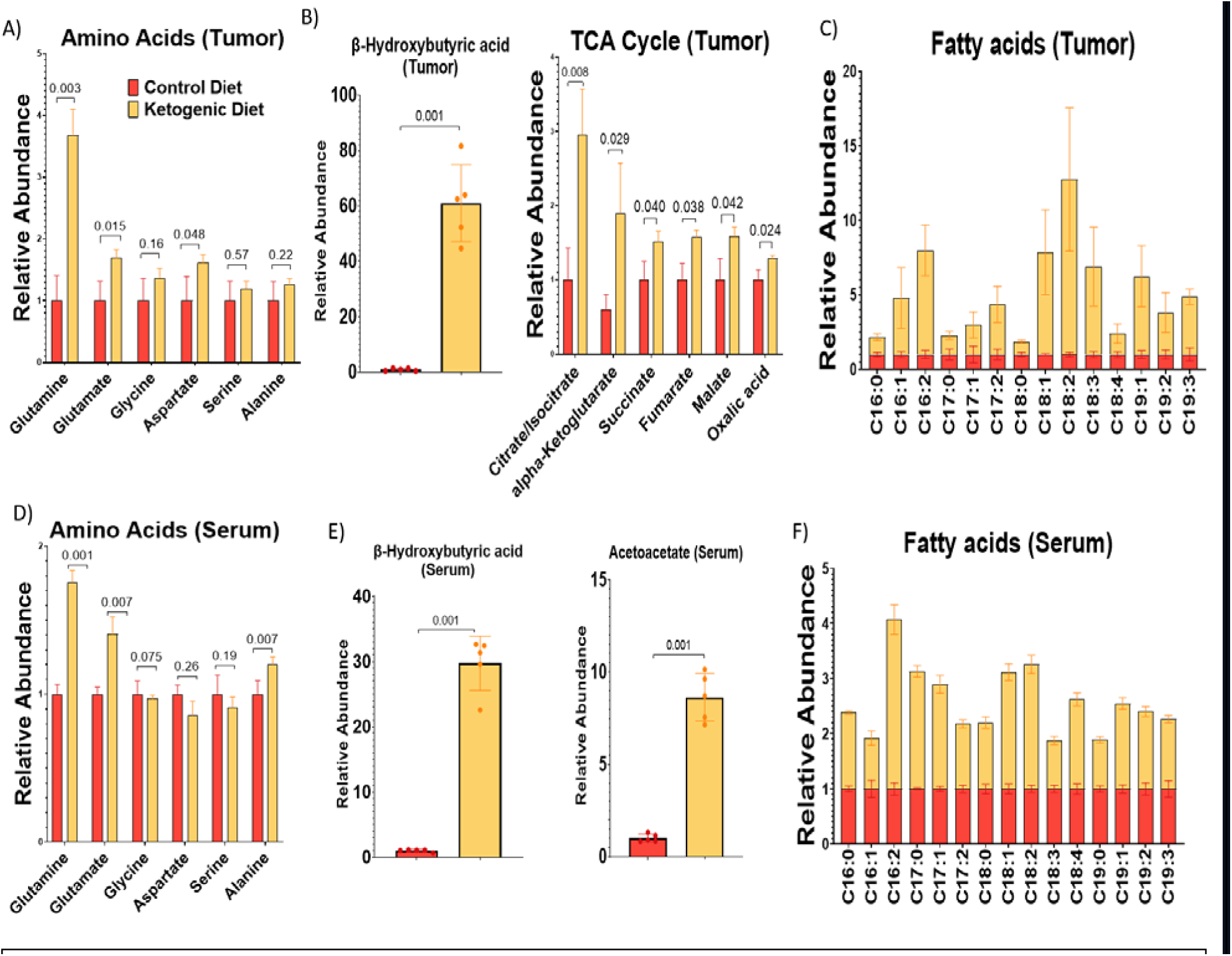
Impact of a Ketogenic Diet on TCA Cycle and Downstream Metabolites in a PDAC Mouse Model. MIA-PaCa2 tumors were injected into mice, and after 15 days of treatment, the tumors were harvested. **(A)** Relative levels of glutamine, glutamate, glycine, aspartate, serine, and alanine were measured in the tumors of mice. **(B)** Levels of β-hydroxybutyric acid and TCA cycle metabolites in tumors. **(C)** Relative comparison of circulating fatty acids in tumors of mice on a ketogenic diet versus a control diet. **(D)** Serum levels of amino acids. **(E)** Levels of ketone bodies circulating in the serum. **(F)** Comparison of circulating fatty acids in the serum of mice on a ketogenic diet versus a control diet.

We next assessed the impact of glutamine availability on MIA-PaCa2 cells *in vitro*. The results demonstrated that a 50% reduction in glutamine levels led to a corresponding 50% decrease in cell viability, even under high glucose conditions (25mM), which are characteristic of standard cell culture media (Supplementary Data Fig. 2A). Under low glucose conditions (2.5mM, as a simulation of a key effect of a ketogenic diet on glucose levels in circulation and tumors)(26), cell viability was hardly apparent even with modest glutamine reductions (Supplementary Data Fig. 2B). These data indicate that PDAC cells rely on glutamine availability, especially under low glucose conditions.

### Specific nutrient components of a ketogenic diet alter metabolism of PDAC cells

In a series of isotope tracing experiments, we investigated the metabolic contributions of uniformly labeled glucose, caprylic acid, β-hydroxybutyrate, and glutamine to PDAC cell metabolism. Conditions were intentionally modified with relatively low glucose and high fats and ketone bodies to simulate a ketogenic diet. The results indicated that glucose predominantly fuels the glycolysis pathway (e.g., M+3 pyruvate and lactate in Fig. 3A), while caprylic acid, β-hydroxybutyrate, and glutamine are directly integrated into the TCA cycle (Fig. 3A, M+2 TCA metabolites for caprylic acid and BHB; M+4/5 metabolites for glutamine). Mechanistically, caprylic acid and β-hydroxybutyrate enter the TCA cycle via acetyl-CoA, whereas glutamine is metabolized through glutamate and subsequently alpha-ketoglutarate (Fig. 3A & B).

**Figure 3:**
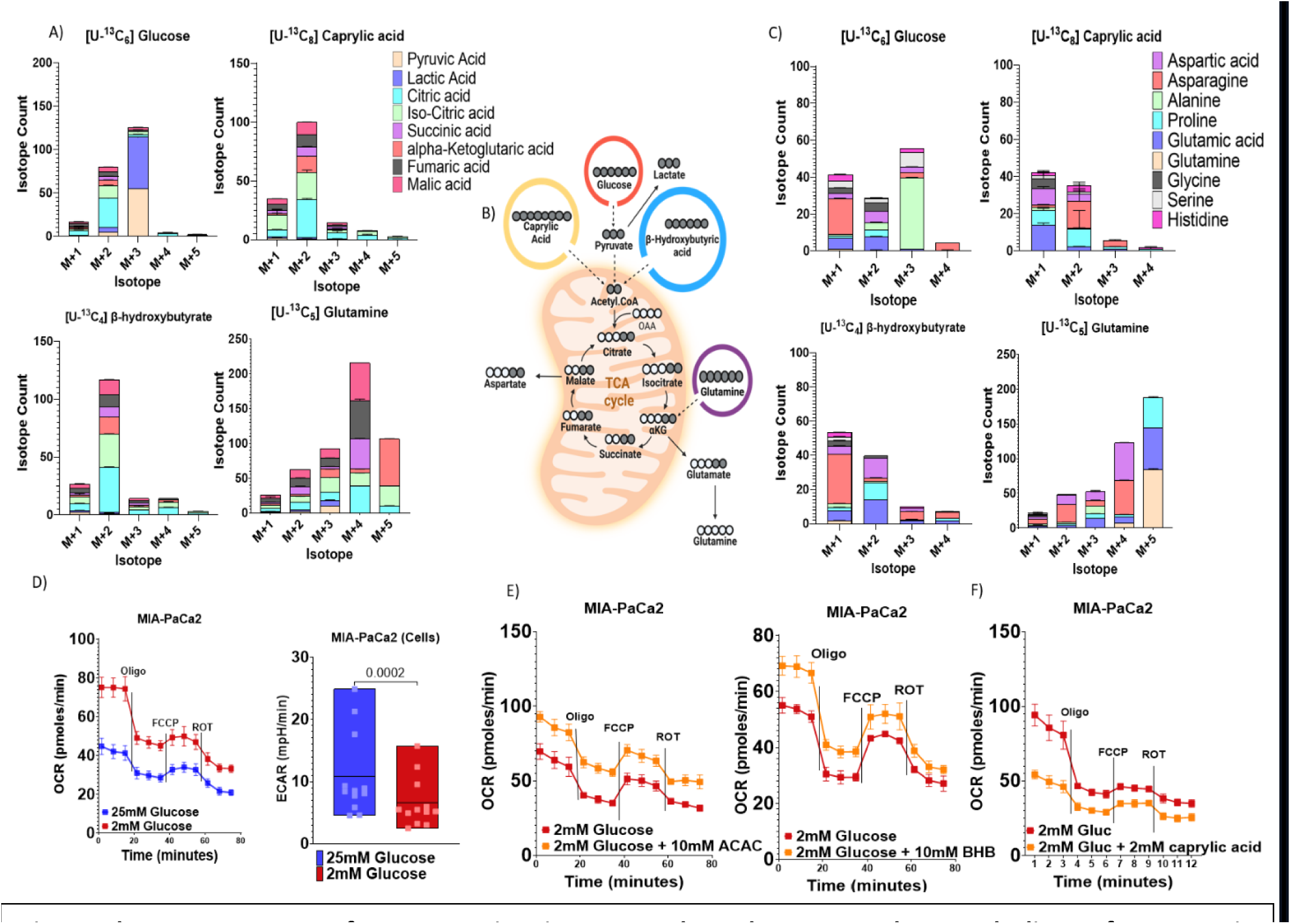
The Components of a Ketogenic Diet Upregulate the TCA Cycle Metabolism of Pancreatic Cancer. The collective pool size of caprylic acid, glutamine, glucose, and β-hydroxybutyric acid uptake was assessed via mass spectrometry in MiaPaCa-2 cells cultured with 4 mM [U-13C] glutamine, 10 mM DL-BHB, 2mM octanoic acid, and 5 mM glucose for 3 hours. **(A)** Contribution of labeled glucose, caprylic acid, β-hydroxybutyric, and glutamine to TCA cycle (n=3 independent biological replicates). **(B)** Schematic to illustrate the circulation of labeled carbon and its contribution to the TCA cycle. **(C)** Isotopologue distribution of labeled carbon distribution to amino acid synthesis. **(D)** Comparison of oxygen consumption rate and extracellular acidification rate between 25mM glucose and 2mM glucose conditions. **(E-F)** OCR activity in MIA-PaCa2 cells treated with low glucose compared to low glucose supplemented with ketone bodies and caprylic acid.

Further analysis focused on identifying which tracers predominantly label amino acids within the cells. The data revealed that glutamine serves as a major precursor for the synthesis of several amino acids, including glutamine, glutamate, aspartate, histidine, and asparagine (Fig 3C). While the other nutrients also contributed, comparison of these panels reveal that the magnitude of the contribution was far greater from glutamine, as measured by total isotope counts and the number of labeled carbons comprising these amino acids. Additionally, a series of Seahorse experiments assessed the impact of these components on mitochondrial activity. The findings demonstrated that low glucose conditions and the presence of ketone bodies enhance mitochondrial activity. Unexpectedly, the treatment of cells with caprylic acid under low glucose conditions resulted in a reduction of mitochondrial activity (Fig. 3D & E). Collectively, these data suggest that PDAC cells undergo a metabolic shift in response to a ketogenic diet, with the diet’s components generally upregulating TCA metabolite levels and mitochondrial activity.

### Ketone bodies and fatty acids are critical fuel sources for the TCA cycle in the context of a ketogenic diet

Data from above were reanalyzed in order to more directly compare the impact of ketogenic diet components on the TCA cycle and cataplerosis (exiting the TCA cycle). We focused specifically on M+2 carbon and selected M+3 labeled metabolites (Fig. 4A&B). Results indicated that TCA metabolites exhibited significant labeling from [U-^13^C_4_]-beta-hydroxybutyrate and [U-^13^C_8_]-caprylic acid (Fig. 4B). Similarly, glutamine proved to be an important anaplerotic substrate for TCA cycle intermediates. Interestingly, labeled carbons from caprylic acid and ketone bodies contributed to the pool of ^13^C-M+2 glutamate and aspartate, presumably through GOT conversion of TCA metabolites, yet no labeled carbons were detected in glutamine from these substrates (Fig. 4C), suggesting minimal net glutamine synthetase activity (which converts glutamate to glutamine).

**Figure 4:**
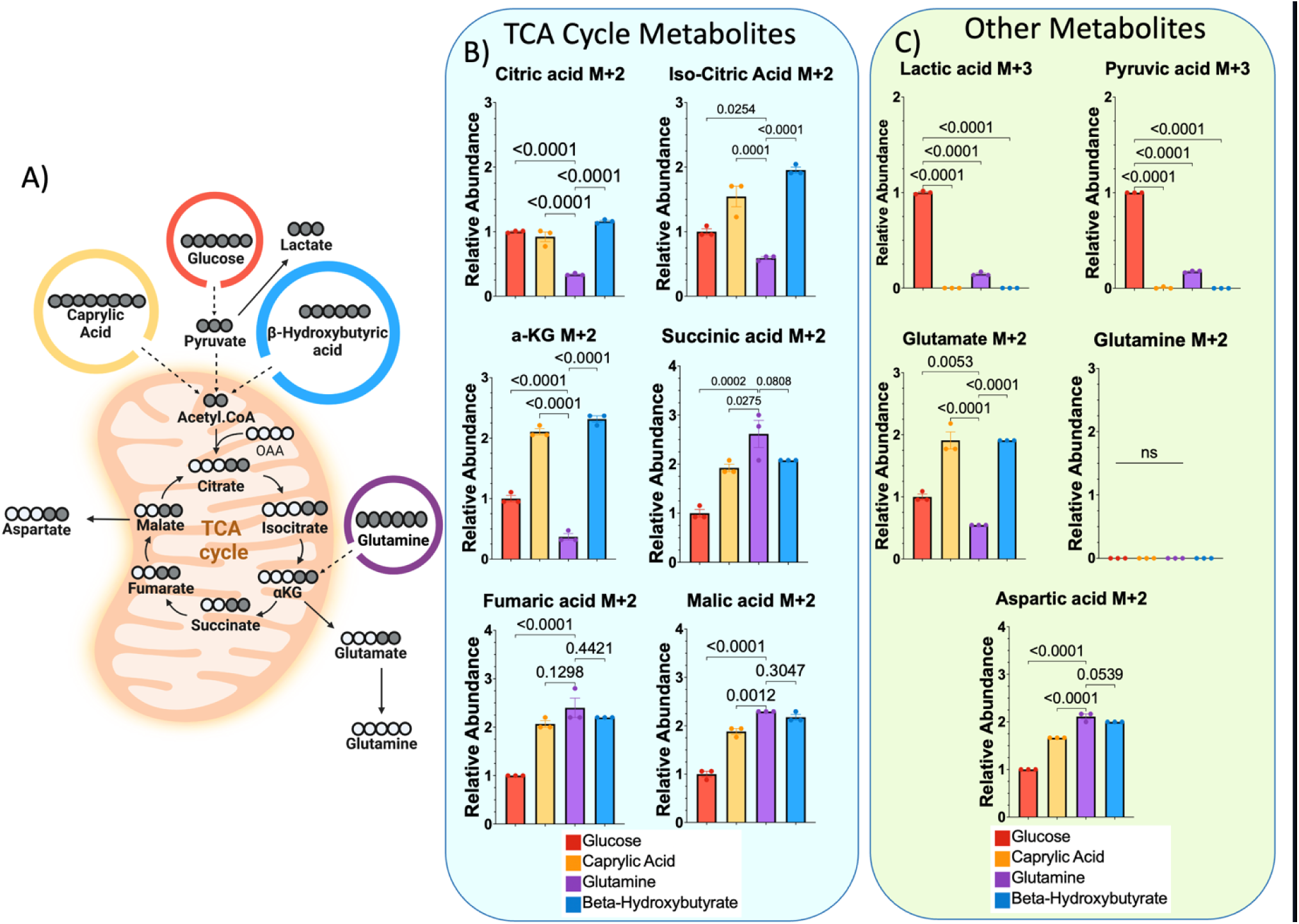
Contribution of Ketogenic Diet Components to Various TCA Cycle and Amino Acid Intermediates. The relative uptake of caprylic acid, glutamine, glucose, and β-hydroxybutyric acid was quantified using mass spectrometry in MIA-PaCa2 cells cultured with 4 mM [U-13C] glutamine, 10 mM DL-BHB, 2 mM octanoic acid, and 5 mM glucose for 3 hours (n = 3 individual biological replicates). **(A)** Schematic illustration of isotope tracing experiment. **(B-C)** isotopologue distribution of M+2 and selected M+3 carbon on the indicated metabolites in cells cultured.

### Glutamine is a critical anaplerotic substrate in PDAC cells under ketogenic diet conditions

Similarly, we reanalyzed the same set of experiments to better characterize the extent of glutamine utilization in these core metabolic pathways. Measurements of M+4 and M+5 labeled metabolites from uniformly (^13^C_5_) labeled glutamine revealed that MIA-PaCa2 cells take-up glutamine from their environment, direct carbon into the TCA cycle, and convert these intermediates into associated amino acids (Fig. 5A). Specifically, carbon incorporation from uniformly ^13^C-labeled glutamine, [U-^13^C_5_]-glutamine, was naturally observed as M+5 glutamine (positive control), as well as M+5 glutamate upon conversion by glutaminase (Fig. 5B). Labeled glutamate then directly entered the TCA cycle, as evidenced by elevated M+4 labeled TCA intermediates (Fig. 5C). High levels of M+5 iso-citrate and citrate suggest some degree of TCA cycle reversal as well (i.e., reductive carboxylation of α-ketoglutarate) (Supplementary data Fig. 3A-B). Labeling from glutamine was observed in other key amino acid intermediates, including proline (all 5 carbons derived from glutamine, hence M+5) and asparagine (synthesis requires NH_3_ donation from glutamine, as well as oxaloacetate for the carbon backbone, hence M+4) (Fig. 5B). An additional round through the TCA cycle yields M+3 TCA cycle intermediates (Supplementary Fig. 3C).

**Figure 5:**
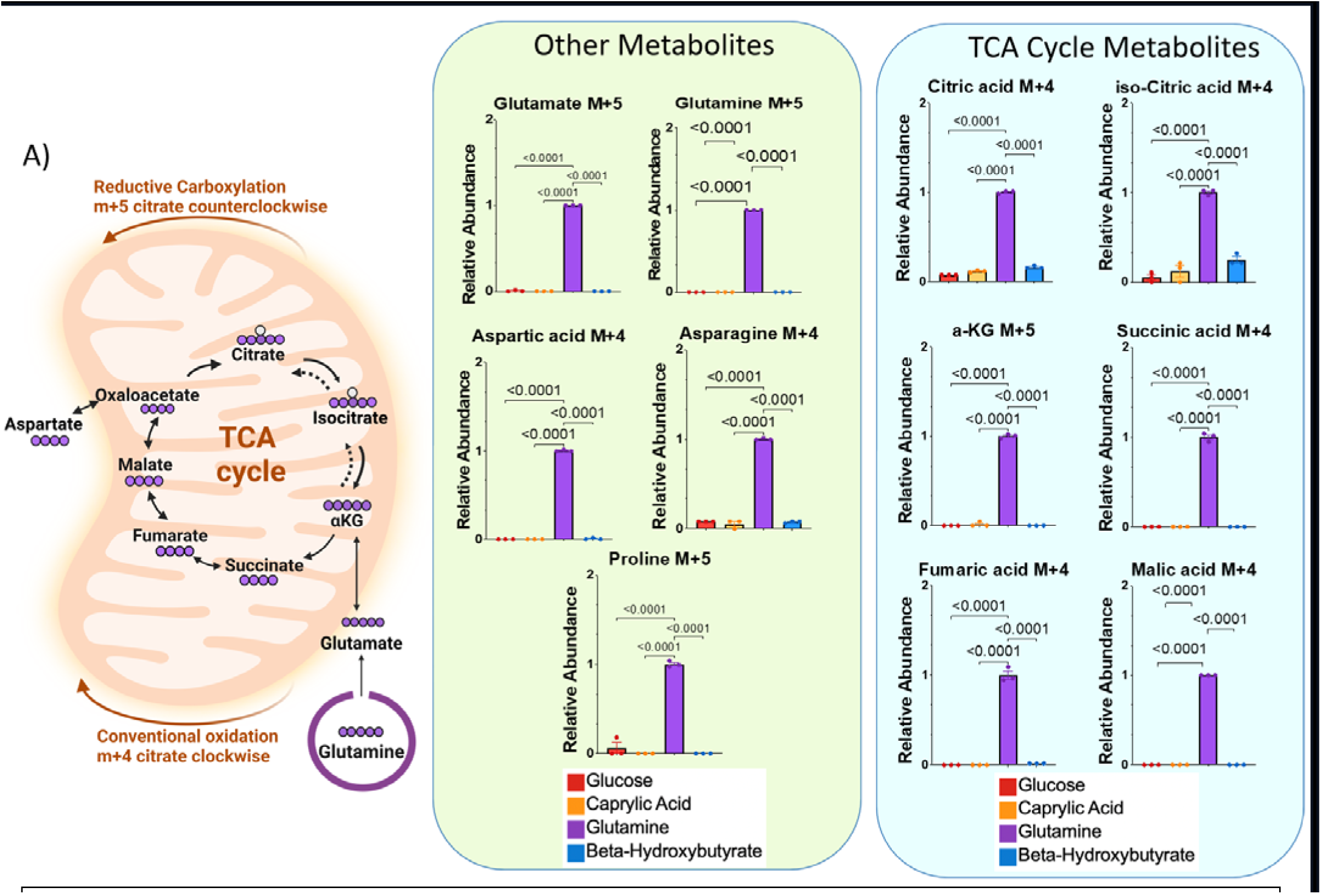
Assessment of the Impact of Glutamine and Other Components of a Ketogenic Diet on the TCA Cycle and Amino Acid Intermediates. **(A)** Schematic to illustrate the circulation of uniformly labeled glutamine and its contribution to the TCA cycle. **(B-C)** The collective pool size of caprylic acid, glutamine, glucose, and β-hydroxybutyric acid uptake was assessed via mass spectrometry in MiaPaCa-2 cells cultured with 4 mM [U-C_M_] glutamine, DL-BHB 10mM, Octanoic acid 2mM, and Glucose 5mM for 3 hours (n = 3 independent biological replicates). **(B-C)** Isotopologue distribution of M+5 carbon on the specified metabolites in cultured cells.

### A ketogenic diet potentiates glutamine metabolism inhibition to treat PDAC

Based on these data in aggregate, we reasoned that PDAC cells become reliant on glutamine in the context of a ketogenic diet and may be vulnerable to glutamine inhibition in combination with the diet. To test this possibility, we performed independent flank xenograft studies using human MIA-PaCa2 cells in nude mice and KPC cells in syngeneic C57BL/6 mice. In the former experiment, the pan-glutamine metabolism inhibitor, DON was used, while a GLS inhibitor was used in the latter experiment. After 10 days of co-treatment, we observed a significant suppression of tumor growth in the combination therapy groups compared to diet or glutamine inhibition alone (Fig. 6A & B). Measurements of β-hydroxybutyrate confirmed the expected metabolic effects of a ketogenic diet (Fig. 6C). Body weights were slightly lower in mice receiving a ketogenic diet plus DON, but were overall stable, indicating the diet plus drug combination was not toxic (Fig 6D). Thus, we propose a model where limited glucose availability in ketogenic diet shifts PDAC metabolism towards reliance on glutamine anaplerosis, and thhs metabolic rewiring sensitizes PDAC tumors to glutamine metabolism inhibition.

**Figure 6:**
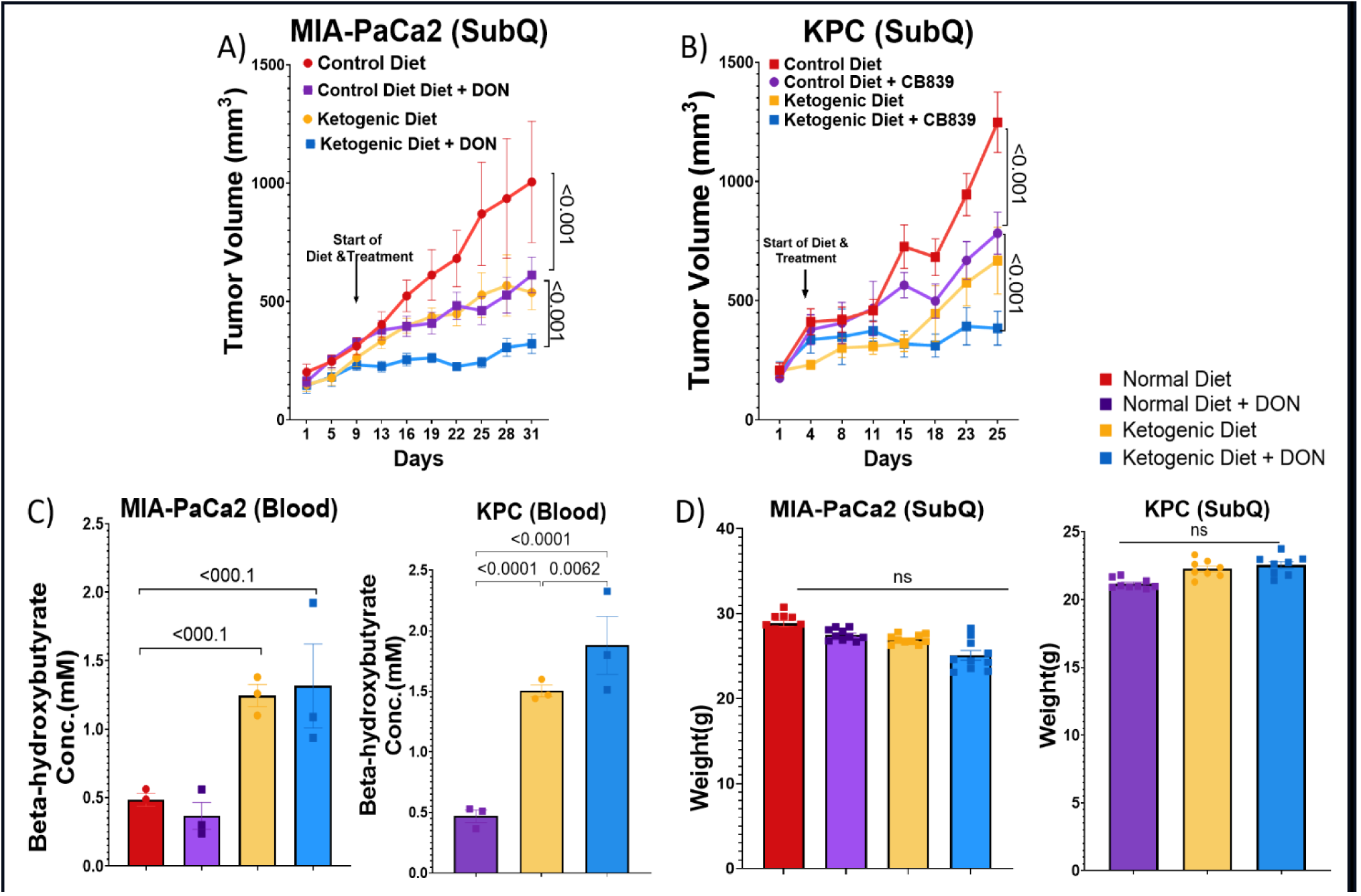
Synergistic Inhibition of Tumor Growth with Ketogenic Diet and Glutamine Metabolism Inhibitors. **(A)** MIA-PaCa2 tumor volumes in nude athymic mice treated with DON in combination of with a ketogenic diet (n=7). **(B)** Subcutaneous KPC tumor volumes in C57BL/6 mice treated with CB839 in combination with a ketogenic diet (n=7). **(C-D)** Measured levels of β -hydroxybutyrate and body weight.

## Discussion

KRAS-driven PDAC tumors (90% of all PDAC) have been shown to depend on glutamine metabolism (26–29), likely due to the fact that glucose levels are already depleted in the PDAC TME(26,30). Studies reveal that KRAS-mutant PDAC tumors upregulate both GOT1 and GOT2 (glutamate oxaloacetate transaminase) to utilize glutamine to sustain the TCA cycle (28). Thus, one would hypothesize that PDAC would be susceptible to glutamine metabolism targeted treatments. Unfortunately, preclinical studies investigating glutamine inhibition as a therapeutic strategy have been equivocal. For instance, Biancur et al. observed that the oral glutaminase inhibitor CB-839 did not affect tumor progression in LSL-KRAS^G12D^;p53^L/+^;PDX1-Cre tumors or human PDAC flank xenografts in nude mice (31). Son et al. found that genetic targeting of the gln metabolism downstream enzymes GOT1 or GOT2, resulted in a significant decrease in tumor volume in a human PDAC xenograft model, indicating these enzymes could serve as therapeutic targets (28). In accordance, Recouvreux et al. reported significant decrease in tumor volume in several PDAC orthotopic models when using the pan-glutamine targeting drug, DON (32). Concerns regarding DON-related toxicity prompted the development of DRP-104, a pro-drug version (32), which showed promising responses in a syngeneic PDAC model (33). Combination therapy with trametinib, a MEK inhibitor, significantly increased survival in the same model, indicating the superior potential of glutamine inhibition when combined with rationally selected complementary therapies (33).

The success of glutamine metabolism inhibition in some preclinical studies spurred the pursuit of developing inhibitors and testing in human clinical trials. As early as 1990, the potential of DON, a pan-glutamine metabolism inhibitor, was explored in late-stage clinical trials in sarcoma and mesothelioma patients. DON demonstrated poor efficacy in clinical trials, with one study of 36 evaluable sarcoma or mesothelioma reporting no objective response(34). To mitigate the toxicity associated with glutamine inhibition, more selective enzyme-targeting drugs such as GLS inhibitors were developed. Telaglenastat (CB-839) has been the most widely studied glutaminase inhibitor, tested in 20 cancer related clinical trials available on clinicaltrials.gov. Of these, 17 trials focused on solid tumors, and findings from 4 trials having been published. A phase 1 clinical trial evaluated CB-839 in addition to standard of care paclitaxel in patients with triple negative breast cancer, with some patients having partial responses despite paclitaxel refractory disease (35). A phase1-2 clinical trial utilizing CB-839 plus capecitabine demonstrated no efficacy by RECIST criteria in solid tumors including the majority of colorectal, cholangiocarcinoma, breast and gall bladder cancers, but a subgroup analysis of PIK3CA mutant colorectal cancer patients vs PIK3CA wild-type colorectal cancer patients demonstrated a non-significantly prolonged progression free survival (24.8 vs 16 weeks, p=0.198). (35,36) These clinical trials are still awaiting the publication of the phase two efficacy results(36,37).

Two published late-phase clinical trials evaluating telaglenastat in combination with other therapies in metastatic renal cell carcinoma provide contradicting results. The phase 2 ENTRATA trial randomized 69 patients to teleglenastat plus evirolimus vs. placebo plus evirolimus. Notably, evirolimus is an mTORC1 inhibitor that disrupts glucose metabolism in cancer cells by reducing glucose uptake, perhaps sensitizing cancer cells to glutamine inhibition (38). Patients receiving the teleglenastat plus evirolimus combination had a small, but non-significant improvement in progression free survival (3.9 months vs 1.9 months p=0.079)(39). The phase 3 CANTANA clinical trial evaluated telaglenstat plus cabozantinib vs. placebo plus cabozantinib in 444 patients with metastatic renal cell carcinoma. Cabozantinib is a MET and VEGFR inhibitor, and has shown similar metabolic effects to evirolimus in cancer cells, namely decreasing glucose consumption(40). There was no difference in the primary endpoint of progression free survival (p=0.65)(41). Thus, the drug has marginal activity in patients, and potential therapeutic combination remains to be identified.

Herein, we show that ketogenic diet enhances glutamine metabolism in PDAC, which could be leveraged into a novel combination therapeutic strategy using a ketogenic diet to drive stronger glutamine dependence. The effect was demonstrated through analysis of metabolites, especially in the TCA cycle, in mice fed a ketogenic diet. Additionally, in vitro isotope tracer studies revealed the utilization of ketogenic diet related fuel sources including glutamine into the TCA cycle. Furthermore, ketone bodies and caprylic acid (a medium-chain fatty acid) demonstrated significant carbon incorporation into the TCA cycle. It is important to note that while this study employed a medium-chain fatty acid, a prolonged ketogenic diet is likely to elevate circulating levels of diverse fatty acids derived from dietary fat sources. Most importantly, we show that ketogenic diet sensitizes tumors to glutamine inhibitors, both DON and CB839.

Future studies by our team will aim to validate increased glutamine utilization under ketogenic conditions through *in vivo* isotopologue analyses. In addition, dependence on mitochondrial metabolism can be accentuated further by adding chemotherapy or targeted agents previously tested in clinical trials(42), offering a triple therapy strategy to sensitize cancer cells. While chemotherapy and CB-839 have not shown substantial efficacy in combination to this point (43,44), including a ketogenic diet may provide a necessary shift in the metabolic program required to see an anti-cancer signal in patients. It will be also important to evaluate other metabolic modulators in combination with a ketogenic diet, including drugs that target glutamine metabolism (JHU083, azaserine, and acivicin) or other enzymes fundamental to core metabolic pathways.

## Methods

### Cell lines, cell culture, and reagents

The human pancreatic cancer cell line MIA PaCa-2 was obtained from the American Type Culture Collection (ATCC) (no. CRL-1420). The murine pancreatic cancer cell line (KPC K8484: Kras^G12D/+^; Trp53^R172H/+^; Pdx1-Cre) was provided by the Darren Carpizo laboratory. These two cell lines were maintained at conditions of 37°C and 5% CO_2_. All cells were cultured in DMEM containing 4 mM glutamine and 25 mM glucose, supplemented with 1% penicillin/streptomycin, 10% FBS, and prophylactic doses of plasmocin (Life Technologies, no. MPP-01-03) to prevent mycoplasma infection. A MycoAlert detection kit (Lonza) was subsequently utilized for Mycoplasma screening. To simulate low-glucose conditions in a pancreatic cancer microenvironment, glucose withdrawal was performed as indicated. For low-glucose experiments, glucose-free DMEM (Life Technologies, no. 21013-024) was supplemented with 10% FBS and penicillin/streptomycin, and 2.5 mM glucose.

### Clonogenic assay

Cells were plated in six-well plates at 1,500 cells per well. cells were first cultured with media for 24 h, followed by treatment under varying levels of glutamine, 10% FBS, and the indicated glucose concentrations. At the conclusion of experiments, colonies were fixed in a reagent containing 80% methanol and stained with 0.5% crystal violet. To determine relative growth, dye was withdrawn from stained colonies with 10% acetic acid and the associated absorbance was measured using a microplate reader at 600 nm (GloMax Explorer system, Promega)(45).

### In vivo-studies

All experiments involving mice usage were conducted under the approval of Case Western Reserve University Institutional Animal Care Regulations and Use Committee (CWRU; IACUC protocol no. 2018-0063). Six-to-eight-week-old, female, athymic nude mice (Foxn1 nu/nu) were purchased from Harlan Laboratories (no. 6903M) through ARC CWRU. Mice were sustained in the humanity-controlled animal facility with standard chow (LabDiet, Prolab IsoPro RMH3000), ALPHA-dri bedding (nutrient-free), and under pathogen-free conditions. No additional nutrient-contained bedding or food was provided to the animals during these studies. Mice were fed ad libitum.

For xenograft experiments, MIA PaCa-2cells were suspended in 200μL of a PBS:Matrigel solution (1:1). 1×10^6^ suspended cells were injected subcutaneously into the right flank of mice. The KPC allograft tumor study was conducted in eight-week-old syngeneic female C57BL/6 mice(46), and mice were injected subcutaneously with 5×10^4^ KPC Matrigel suspended cells in the same manner into the right flank. In orthotopic syngeneic experiments, a suspension of 1:1 Matrigel with PBS of 5×10^4^ KPC cells expressing Luciferase was injected directly into the mouse pancreas. On the 10th day after the surgery, the presence of pancreatic tumors was confirmed using bioluminescence imaging (BLI) via Spectrum CT (PerkinElmer, 2898979) after injecting 100 μL D-luciferin (50 mg/mL in PBS) intraperitoneally.

In the therapeutic experiments, treatments started once the tumors were palpable, with tumor volumes averaging 120–150 mm^3^ across treatment groups (47,48). KPC tumors in C57BL/6J mice (n = 36) were randomized to the following treatment arms: Normal diet (ND) (n = 9), Ketogenic diet (KD) (n = 10), ND + CB-839 (n = 8), KD + CB-839 (n = 9). Treatments were administered after two initial tumor measurements. CB-839 was dissolved in a 3 mL solution consisting of 2 mL of vehicle (sterilized H_2_O, NaCl, polyethylene glycol, Tween 80) and 1 mL of corn oil. Both the control and CB-839 (200 mg/kg) treatment group mice were treated via oral gavage three times per week. Experiments testing DON were pereformed in nude mice, with groups randomized to normal diet (ND; BioServ, no. F3155) + vehicle (sterilized H_2_O, n=5), ketogenic diet + vehicle (KD; BioServ, no. F3666, n=5), ND + DON (n=8), and KD + DON (n=9). After confirming the tumor progression by two initial tumor measurement treatments were administered 2 mg of DON was suspended in 4 mL of vehicle for the dissolution of DON. Both vehicle (control) and DON treatments were administered interperitoneally (IP) twice per week at a dose of 5 mg(kg^-1^), unless indicated. Upon finalization of animal experiments, mice were euthanized using isoflurane inhalation followed by cervical dislocation. Tumor volume was assessed for each mouse and plotted longitudinally.

Tumor volumes were measured twice per week using a caliper (volume = length x width2/2); body weights measured twice per week. Serum ketone and glucose levels were measured once per week via tail vein sampling using the Precision Xtra Glucose and Ketone Monitoring System from Abbott (no. 98814-65), as well as the Precision Xtra Blood Glucose and Ketone Test Strips from Abbott (no. 9972865; no. 7074565).

### Metabolite Extraction and Derivatization

Cells were treated with 2mM caprylic acid 10mM CIL CAS#23G-0425, (+)-Sodium 3-hydroxybutyrate (CIL CAS# 2483735-72-2), 5mM glucose CIL CAS# CLM-1396-0 and 4 Mm of glutamine (CIL CAS# CLM1822-H), all with fully labeled carbons for 3 hours. Culture media was aspirated from each well of a 6-well plate, and the cells were gently wash with 2 mL of saline solution. The plate was subsequently positioned carefully on ice. A lysis solution consisting of ice-cold methanol, ice-cold water, and 1 mM tricarballylic acid in a 40:20:1 ratio (600 µL per well) was added. Cells were then delicately scraped using a cell scraper while maintaining the plate on ice. The cell suspension underwent vortexing for 10 seconds and centrifugation at 14,000 × g for 10 minutes at 4. The resulting supernatant was combined with 300 µL of chloroform, vigorously vortexed for 30 seconds, and centrifuged at 3,000 × g for 3 minutes at 4. The upper polar phase was cautiously collected and subjected to drying in a SpeedVac. Each sample underwent derivatization with 20 µL of 4% methoxyamine-hydrochloride in pyridine, followed by a 30-minute incubation period at 45. Subsequently, the samples were further derivatized with 25 µL of mtBSTFA + 1% t-BDMCS and incubated for 60 minutes at 45. After centrifugation at 14,000 × g for 10 minutes at 4, the supernatant was collected and processed for gas-chromatography mass spectrometry (GC-MS) analysis.

Metabolites were measured using GC-MS with an Agilent 5977B system, containing an HP-5 ms column (30 m x 0.25 mm, 0.25 µm). The injector temperature was maintained at 300 °C, and 1 µL of each sample was injected. The GC temperature program started at 60 °C, held for 1 minute, increased by 6.5 °C/min to 325 °C, and maintained at 325 °C for 10 minutes. Helium served as the carrier gas with a flow rate of 1.2 mL/min. The analytes underwent electron impact ionization (EI), and metabolite ions were monitored using selected ion monitoring (SIM) mode. The MS source and quadrupole temperature were set at 280 and 150 °C, respectively. The MassHunter software facilitated the annotation of metabolites, providing chromatographic peak areas for the monitored isotopomer peaks. Subsequently, IsoCorrectoR was employed to correct for natural isotope abundance and derive the isotopomer distribution for each metabolite.

LC-MS analysis was used to measure water-soluble metabolits by running samples on the Orbitrap Exploris 480 mass spectrometer (Thermo Scientific) coupled with hydrophilic interaction chromatography (HILIC) and an XBridge BEH Amide column (150mm X 2.1 mm, 2.5 uM particle size, Waters, Milford, MA). The gradient included solvent A (95%:5% H2O:acetonitrile with 20 mM ammonium acetate, 20 mM ammonium hydroxide, pH 9.4) and solvent B (100% acetonitrile) according to the following times and ratios: 0min, 90% B; 2min, 90% B; 3min, 75% B; 7min, 75% B; 8min, 70% B, 9min, 70% B; 10 min, 50% B; 12 min, 50% B; 13 min, 25% B; 14 min, 25% B; 16 min, 0.5% B, 20.5 min, 0.5% B; 21 min, 90% B; and 25 min, 90% B. The flow rate was 150 uL/min with an injection volume of 5 uL and a column temperature of 25 °C. The MS scans were in polarity switching mode to enable both positive and negative ions across a mass range of 70–1000 m/z, with a resolution of 120,000. Data were analyzed using the EI-MAVEN software (v 0.12.0, Elucidata).

Statistical analyses. *In vitro* data are presented as mean ± s.e.m. of at least three independent experiments unless indicated. Comparisons between groups were determined using the unpaired, two-tailed Student *t-*test (* *p* < 0.05; ** *p* < 0.01; *** *p* < 0.001 **** *p* < 0.0001). One-way or Two-way ANOVA tests were used to compare more than two groups. Kaplan-Meier estimation was used with the log-rank tests to estimate distributions of survival (19) GraphPad Prism 9.2.3 Software was used for statistical analyses.

**Table 1:**
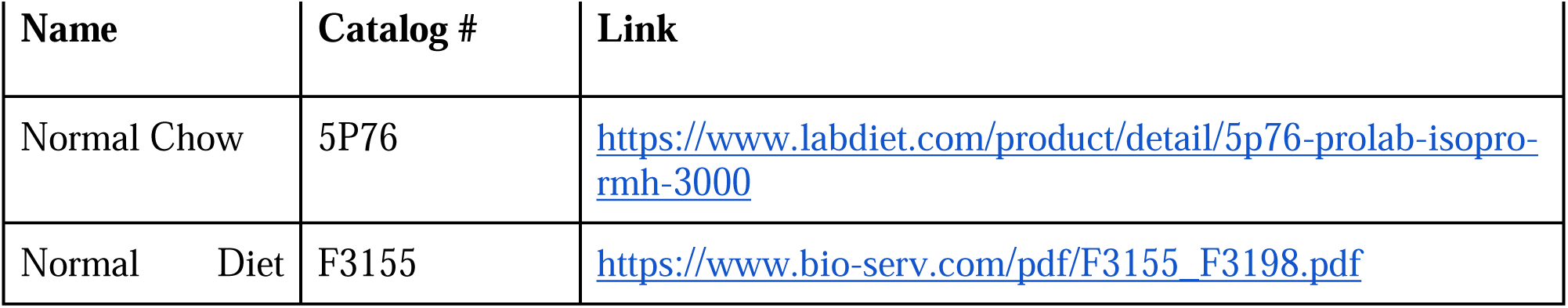
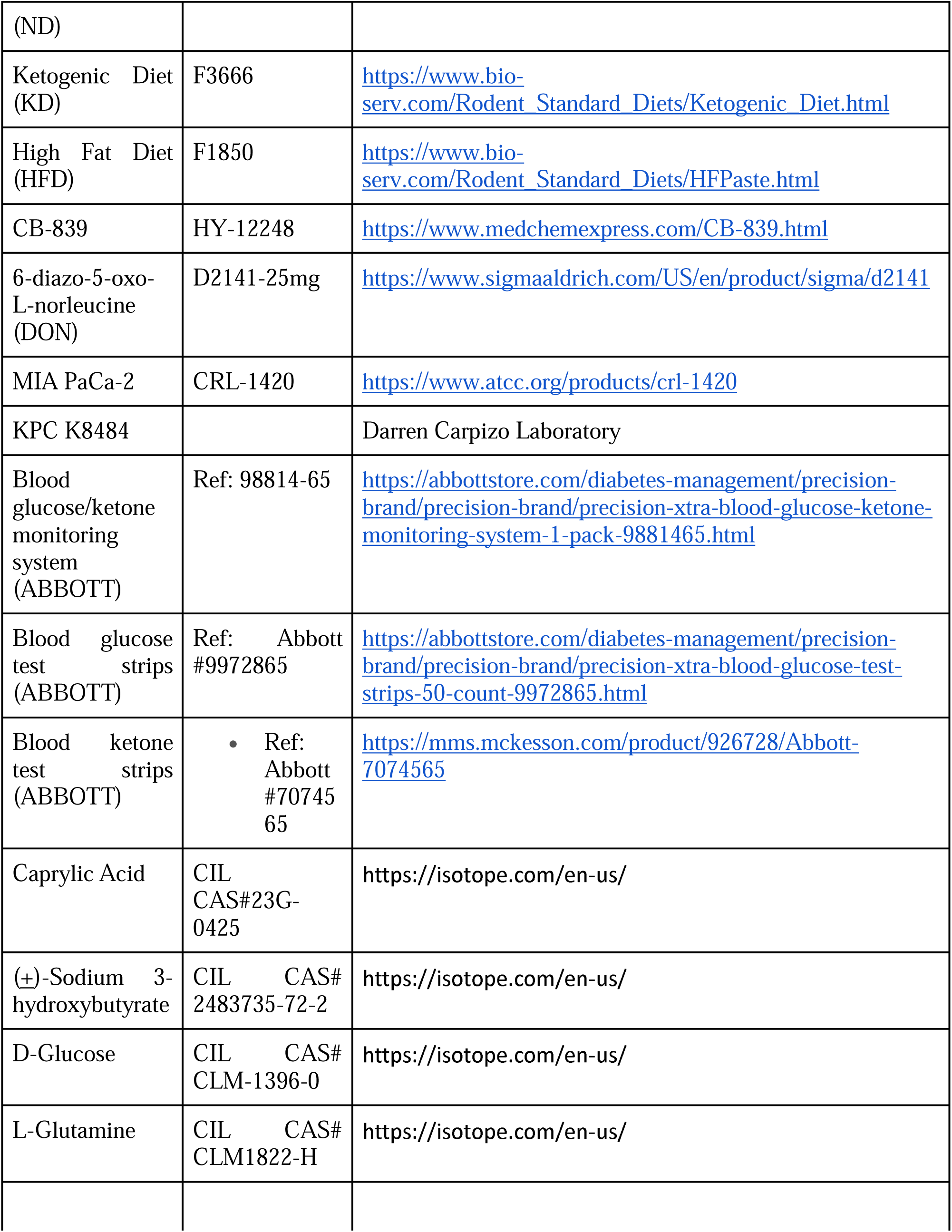
List of diet and other reagents used in this study

## Disclaimers

We have no disclaimers

---

**Supplementary Fig. 1:**
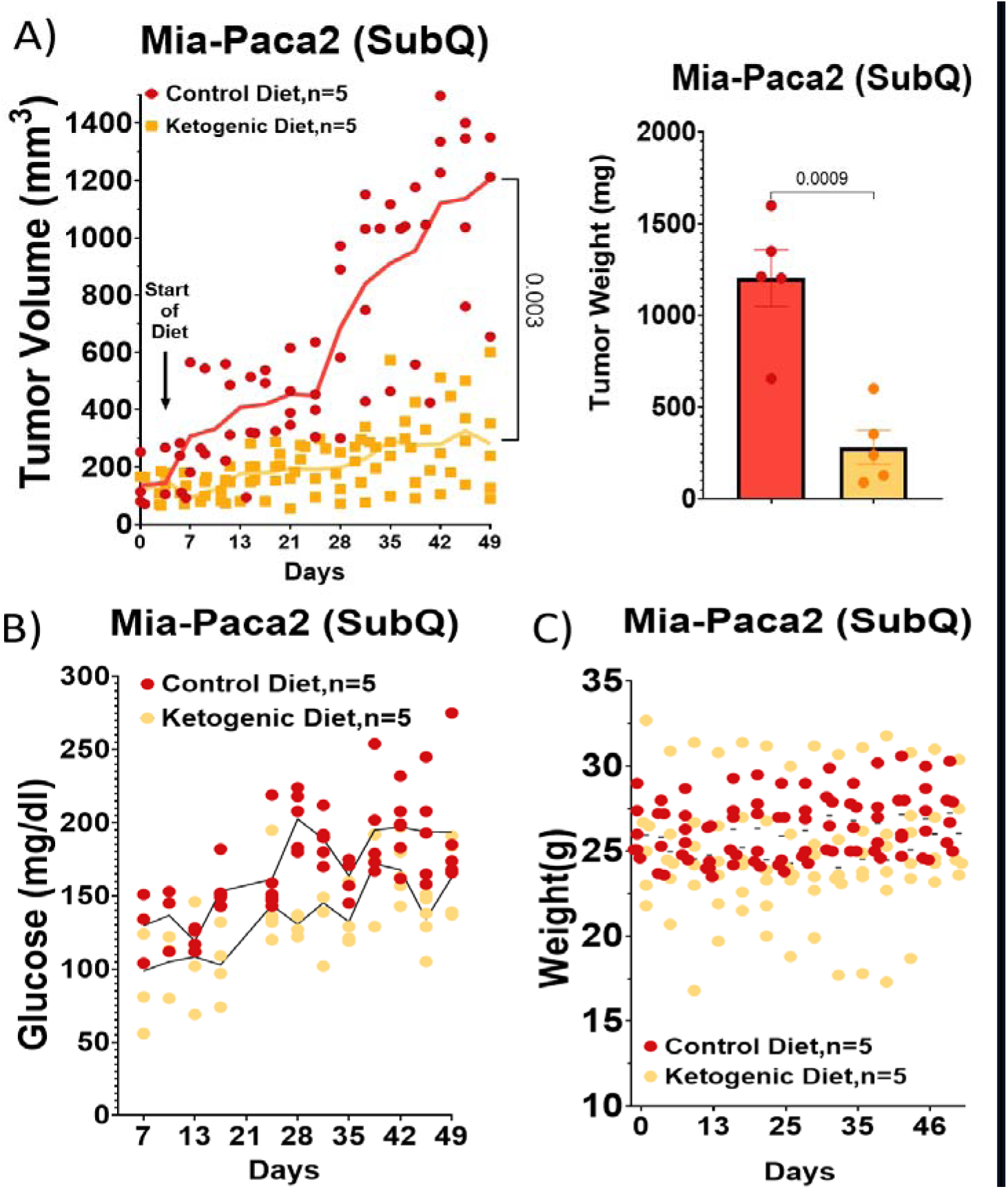
Mice Maintained on a Ketogenic Diet Sustain Lower Blood Glucose Levels While Effectively Managing Their Body Weight. **(A)** MIA-PaCa2 tumors injected in the flank of nude athymic mice. Tumor volume per mouse is illustrated throughout the duration of the experiment, with each day’s measurement representing n=5. Tumors were harvested and weighed upon the completion of the experiment and weighted. **(B)** Glucose levels were measured in individual mice throughout the experiment. **(C)** Mouse body weights were measured twice per week.

**Supplementary Fig. 2:**
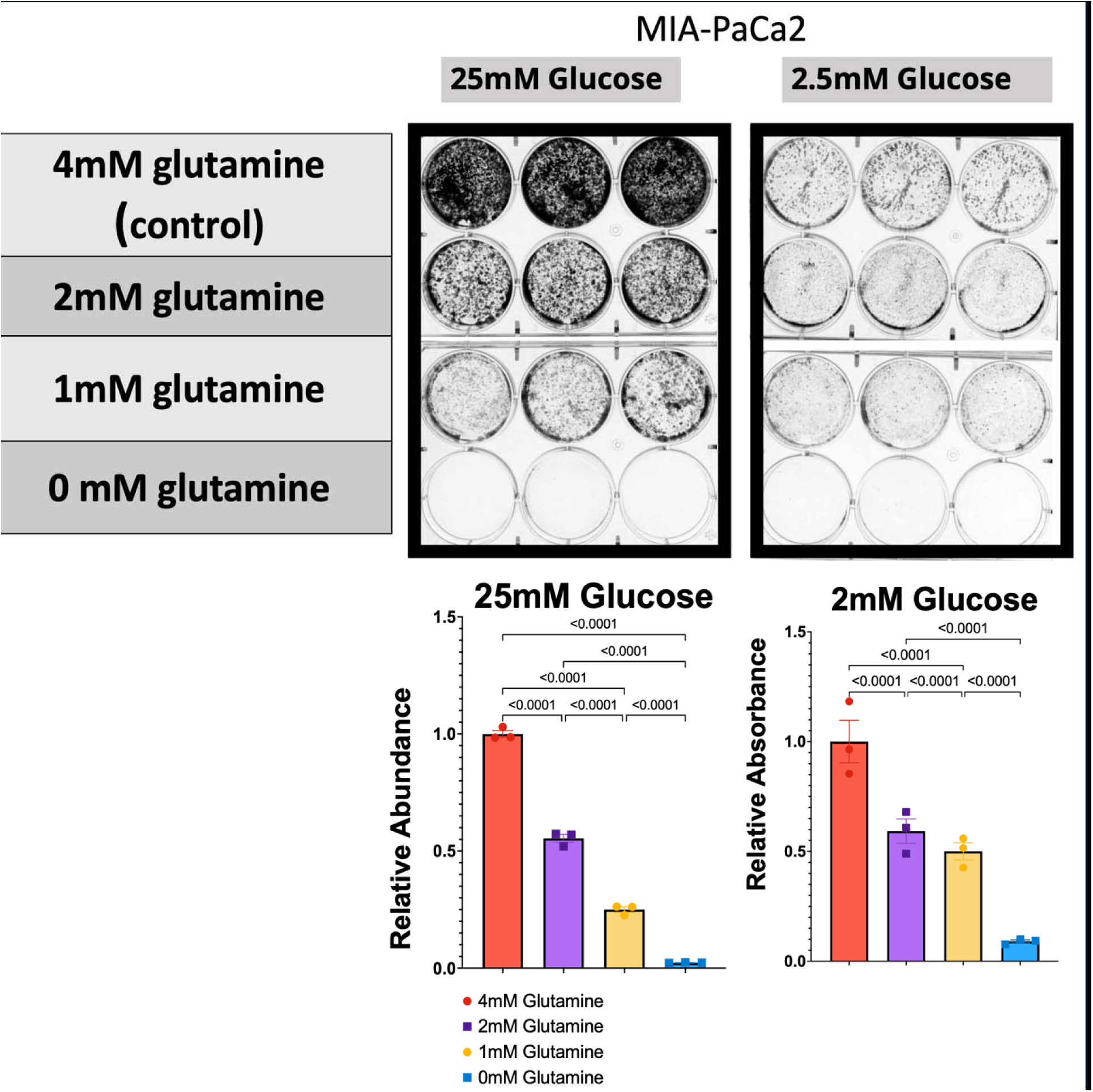
Glutamine is required for Pancreatic Cancer Cells Viability. **(A)** Different glutamine levels were tested under high (25mM) and low (2.5mM) glucose conditions. **(B)** Quantification of cell viability at the different indicated conditions.

**Supplementary Fig. 3:**
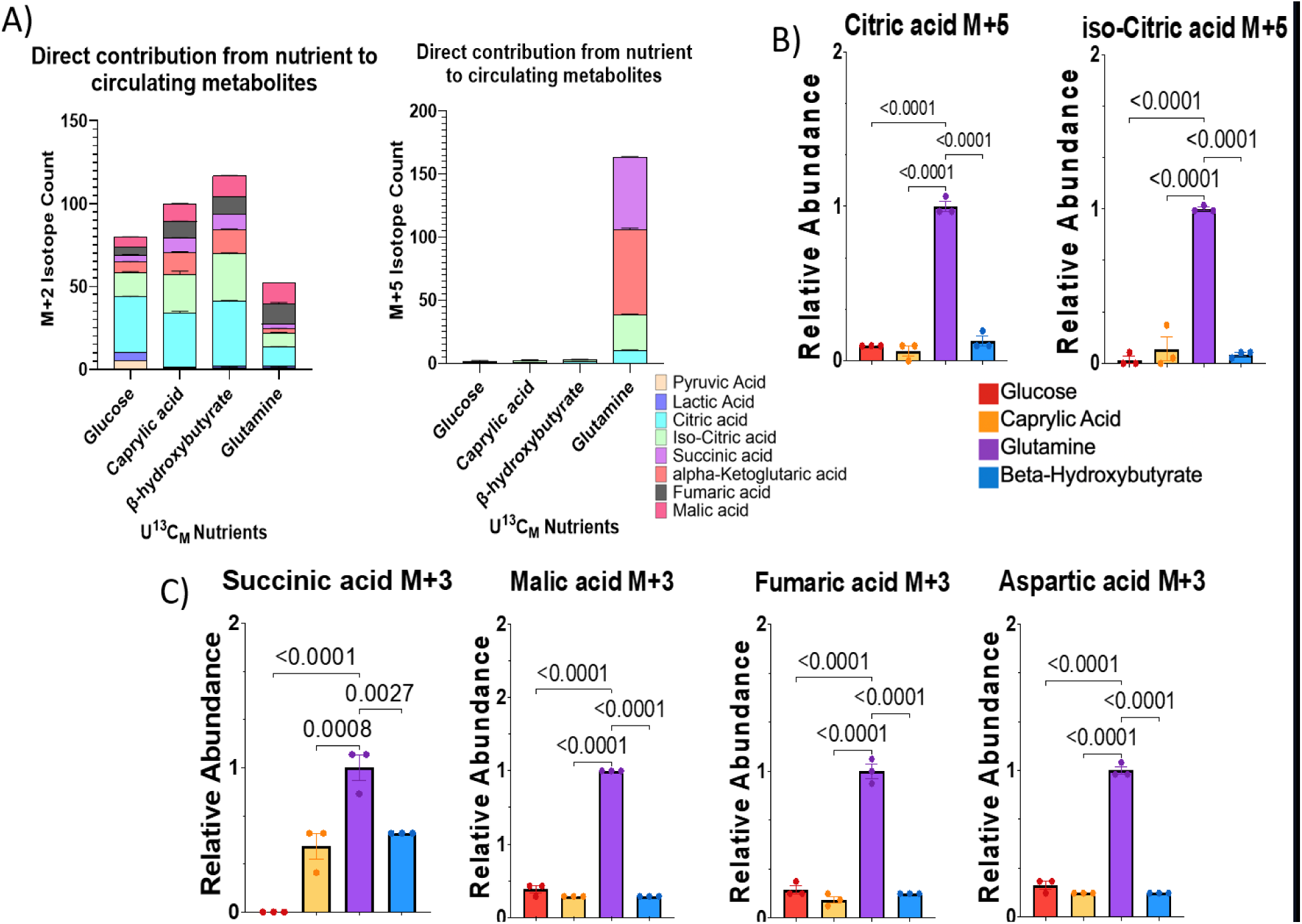
Contribution of Glutamine to the TCA Cycle through Anaplerotic Processes. **(A)** Contribution of ketogenic diet components to M+2 and M+5 of TCA cycle metabolites. **(B)** Contribution of glutamine to the pool of M+5 citric acid and iso-citric acid illustrates a counter-clockwise TCA cycle. **(C)** M+3 pool of succinic acid, malic acid, fumaric acid, and aspartic acid suggests the loss of labeled carbon from M+5 α-ketoglutarate with subsequent rounds through the TCA cycle, while the labeling pattern is attributable to the gain of labeled carbon from the non-labeled-glutamine nutrient sources (glucose, caprylic acid, and BHB).

